# Differential metabolic adaptations define responses of winner and loser oncogenic mutant stem cells in skin epidermis *in vivo*

**DOI:** 10.1101/2022.11.21.517380

**Authors:** Anupama Hemalatha, Zongyu Li, Karen Tai, David G. Gonzalez, Elizabeth Lathrop, Daniel Gil, Catherine Matte-Martone, Smirthy Ganesan, Sangwon Yun, Lauren E. Gonzalez, Melissa Skala, Rachel J. Perry, Valentina Greco

## Abstract

Skin epithelial stem cells detect and correct aberrancies induced by oncogenic mutations. Different oncogenes invoke different mechanisms of epithelial tolerance: while wild-type cells outcompete β-catenin-Gain-of-Function (βcatGOF) mutant cells, Hras^G12V^ mutant cells outcompete wild-type cells^1,2^. Here we ask how metabolic states change as wild-type stem cells interface with mutant cells, and how this ultimately drives different cell competition outcomes. By adapting our live-imaging platform to track endogenous redox ratio (NAD(P)H/FAD) with single cell resolution in the same mice over time, we show that wild-type epidermal stem cells maintain robust redox ratio despite their heterogeneous cell cycle states. We discover that both βcatGOF and Hras^G12V^ models lead to a rapid drop in redox ratios. However, the “winner” cells in each model (wild-type in βcatGOF and mutant in Hras^G12V)^, rapidly recover their redox ratios, irrespective of the mutation induced. Using mass spectrometry (^13^C-LC-MS/MS)^3^, we find that both mutants increase flux through the oxidative tricarboxylic acid cycle, but the “winner” Hras^G12V^ cells and the “loser” βcatGOF cells modulate glycolytic flux differently. Hence, we reveal the metabolic adaptations that define the hallmarks of winners and losers during cell competition *in vivo* and uncover the nodes of regulation unique to each cell fate.

## Introduction

Phenotypically normal human and mouse tissues contain clones of oncogenic mutations, sometimes in frequencies similar to cancer ^4–7^. It is critical to elucidate the cellular mechanisms that maintain tissue function and homeostasis in the presence of oncogenic mutant cells with different growth rates, so that we can understand how those mechanisms are lost in disease and could be restored therapeutically. Using our ability to track oncogenic mutant cells within tissues over time, we have previously shown that genetically mosaic mouse skin cells can use different strategies to correct phenotypic aberrancies caused by oncogenic mutations and to restore function ^1,2,8^. Specifically, while activated β-catenin (βcatGOF) mutant cells are eliminated from the tissue through selective differentiation^1^, constitutively active Hras (Hras^G12V^) mutant cells outcompete wild-type (WT) cells and yet are integrated into normal tissue architecture and function^1,2^. These two mutants and their behaviors in the adult mouse skin invokes cell competition models, with activated βcatGOF cells taking on a “loser” fate and Hras^G12V^ taking on a “winner” fate in comparison to WT cells. Understanding the early responses to either mutation would help uncover the sequence of events that culminate in a cell type being integrated in or being eliminated from the stem cell compartment.

Cell competition, the phenomenon by which unfit cells are eliminated from the tissue by their neighbors, is essential during development, maintenance of organ size, and tumor surveillance ^9,10^. Unfit cells can be removed by apoptosis, apical extrusion, and differentiation, depending on the endogenous cellular behaviors unique to the tissue; in the skin epithelium, the elimination of “unfit” cells happens through selective differentiation.^1,11^. However, the criteria that determine “fitness” of a cell are not well understood. Several signaling pathways including the Wnt and Ras pathways (related to mutations used in this study) have been implicated in this process ^10^. Cell competition is thought to act through modulation of mechanical forces ^12^, apico-basal polarity ^12,13^, and metabolism ^14,15^. Among these, cellular metabolism is sensitive to multiple cues including other sensors of cell competition, like growth factor signaling ^16^ and mechanical forces ^17^, and could potentially integrate upstream cues to directly modulate behaviors involved in tissue growth, such as proliferation ^16,18^ and increase in cell size ^19,20^. Proliferative cells including stem cells disproportionately upregulate glycolysis over mitochondrial oxidation, even in the presence of oxygen, despite glycolysis being far less efficient in generating ATP; this is historically described as the Warburg effect initially observed and most commonly studied in tumors ^21,22^. Hence, understanding the dynamics of metabolic state in stem cell behavior of the highly regenerative skin epidermis in the presence of oncogenic mutations is especially relevant.

It remains difficult to measure metabolic state dynamically throughout time and with intra-cellular resolution while preserving the morphology and integrity of the observed tissue. To directly interrogate the cellular properties that determine cell competition outcomes, we combine live imaging of the mutant mosaic epidermis with Optical Redox imaging ^23–25^ so as to simultaneously determine intra-cellular metabolic rates as the tissue adapts to oncogenic mutations. Revisiting the same epidermal cells in the live animal over long periods of time has previously enabled us to uncover inter-cellular coordination and dynamic behaviors within the epidermal stem cell compartment ^1,26,27^. Tracking the metabolic state of the same epidermal stem cells in the live animal allows us for the first time to bridge the gap between cell behaviors and intra-cellular pathways that define the ability of cells to persist within the tissue.

Optical redox imaging captures the endogenous fluorescence of the reduced NAD(P)H and oxidized FAD such that their relative ratios reflect the balance between reduced and oxidized reactions within the cell ^23–25^. NADH and FAD co-enzymes act in glucose metabolism and their relative ratio represents the balance of glycolytic rate (measured by the reduced NADH levels) and mitochondrial metabolism (measured by oxidized FAD levels) within the cell ^28,29^. The fluorescence detected from the reduced metabolites through optical redox imaging captures both NADH and NADPH (hence called NAD(P)H). However, the ratios of reduced versus oxidized NADP (NADP+ and NADPH), NAD (NAD+ and NADH), and FAD (FAD+ and FADH_2_) are co-dependent and reinforced by one another ^30^ such that NAD(P)H/FAD ratio reflects the redox state of a cell, responsive to glucose catabolic rates. Changes in differential mitochondrial oxidation rates compared to cytosolic glycolysis rates (as expected for oncogenic mutants) will thus be reported by the cumulative optical redox ratio NAD(P)H/FAD ^31,32^.

Using label-free imaging of the endogenous co-enzymes NAD(P)H and FAD, we characterize redox dynamics for the first time in the skin stem cell compartment, and show that NAD(P)H/ FAD is maintained in a robust range distinct from other cell types in the skin, despite variations contributed by changing cell cycle states. Upon encountering the oncogenic mutation βcatGOF, there is a rapid reduction of the NAD(P)H/FAD intensities in the βcatGOF cells and wild-type neighbors at early time points before the appearance of morphological and behavioral aberrancy, making this one of the first observable responses to the mutation *in vivo*. Leveraging revisits over time, we uncover the different redox ratio dynamics in the βcatGOF mosaic stem cell layer as the wild-type cells outcompete the mutant cells to acquire “winner” fate. We find that wild-type cells but not βcatGOF cells recover from the initial drop in NAD(P)H/FAD ratios, leading to a widening redox differential between the two populations. Using a contrasting model Hras^G12V^, wherein the mutant cells outcompete WT cells, we interrogate whether redox changes were mutation-specific or indicative of more general cellular properties like fitness. Strikingly, although there was an initial drop in the NAD(P)H/ FAD intensities, the Hras^G12V^ mutant cells recover their redox ratios with respect to their wild-type neighbors, thereby flattening the redox differential between the two populations over time. Thus, cells that acquire “winner” fate in both oncogenic models recover their redox values, irrespective of the mutation. To identify and define the steps of glucose metabolism (glycolysis, oxidation, carbon exchange) that fuel the changes in redox ratio we observed through imaging in the skin epidermis, we tailored an established liquid chromatography-mass spectrometry/mass spectrometry (LC-MS/MS)-based stable isotope tracer technique ^3^ in awake mice. Using this comprehensive approach to assess key fluxes in cytosolic and mitochondrial glucose metabolism, we discover that the initial changes to redox ratio are caused by an accelerated flux through the TCA cycle in both βcatGOF and Hras^G12V^ mutant epidermis, in line with the reduced NAD(P)H/FAD ratio in both mutants. However, the “winner” mutation Hras^G12V^ specifically upregulates pyruvate to lactate flux and increase *de novo* lactate production while the “loser” βcatGOF mutation downregulates this step. Hence, the winner Hras^G12V^ model differs from the loser βcatGOF mutant model in redox recovery over time as well as glycolytic flux.

Collectively, these metabolic changes are in apparent contrast to the Warburg effect expected for oncogenic cells where the mutant cells are expected to increase glycolytic rates at the expense of TCA cycle and downstream oxidation ^21,22,33^. Instead, in our study, while glucose oxidation is upregulated in both oncogenic mutants, “winner” cells decouple lactate levels (glycolysis) from mitochondrial oxidation; this uncoupling could be a strategy used by “winners” in cell competition. This study also reveals a metabolic plasticity inherent to epidermal stem cells that affect cell competition outcome with implications for therapeutically eliminating oncogenic mutations from the skin epidermis. Thus, we uncover metabolic adaptations *in vivo* that define a winner cell during cell competition in the presence of oncogenic mutations, and show the different metabolic trajectories used to re-establish tissue homeostasis.

## Results

### Optical redox Imaging reveals that epidermal stem cells have a high NAD(P)H to FAD metabolic signature

Metabolic pathways converge at the junction of all major growth factor signals and promote both cell and tissue expansion. The mouse epidermis is constantly renewed by stem cells located in the basal layer (Fig.1A ii), all of which are capable of both proliferating and replenishing the top differentiated cells ^34^. To date, the interplay between the dynamics of metabolism, redox state, and cellular homeostasis *in vivo* is not understood. To address this question, we adapted our previously described two-photon microscopy deep tissue imaging ^34–36^ to incorporate optical redox imaging ^23,24^ and visualize label-free fluorescence of the endogenous metabolites in the hairy (ear) and non-hairy (paw) skin of live mice (Fig1Ai-iii). The intensities captured for redox imaging at specific two-photon excitation wavelengths from intra-vital imaging are collected at wavelength ranges that correspond to NAD(P)H and FAD, and can be represented post-imaging as a ratio NAD(P)H/ FAD (Fig. 1A; Movie S1, S2). To overcome the challenge posed by the undulating three-dimensional tissue in our measurements of NAD(P)H and FAD fluorescence in hundreds of stem cells in the live animal, we isolated the fluorescence signal from the basal layer using distance from the interface of the epidermis and collagen (captured via second harmonic signal) (Fig. S1C-D). Together, these technical advances were critical for measuring endogenous NAD(P)H and FAD intensities in a high-throughput manner and at single-cell resolution.

**Figure 1:**
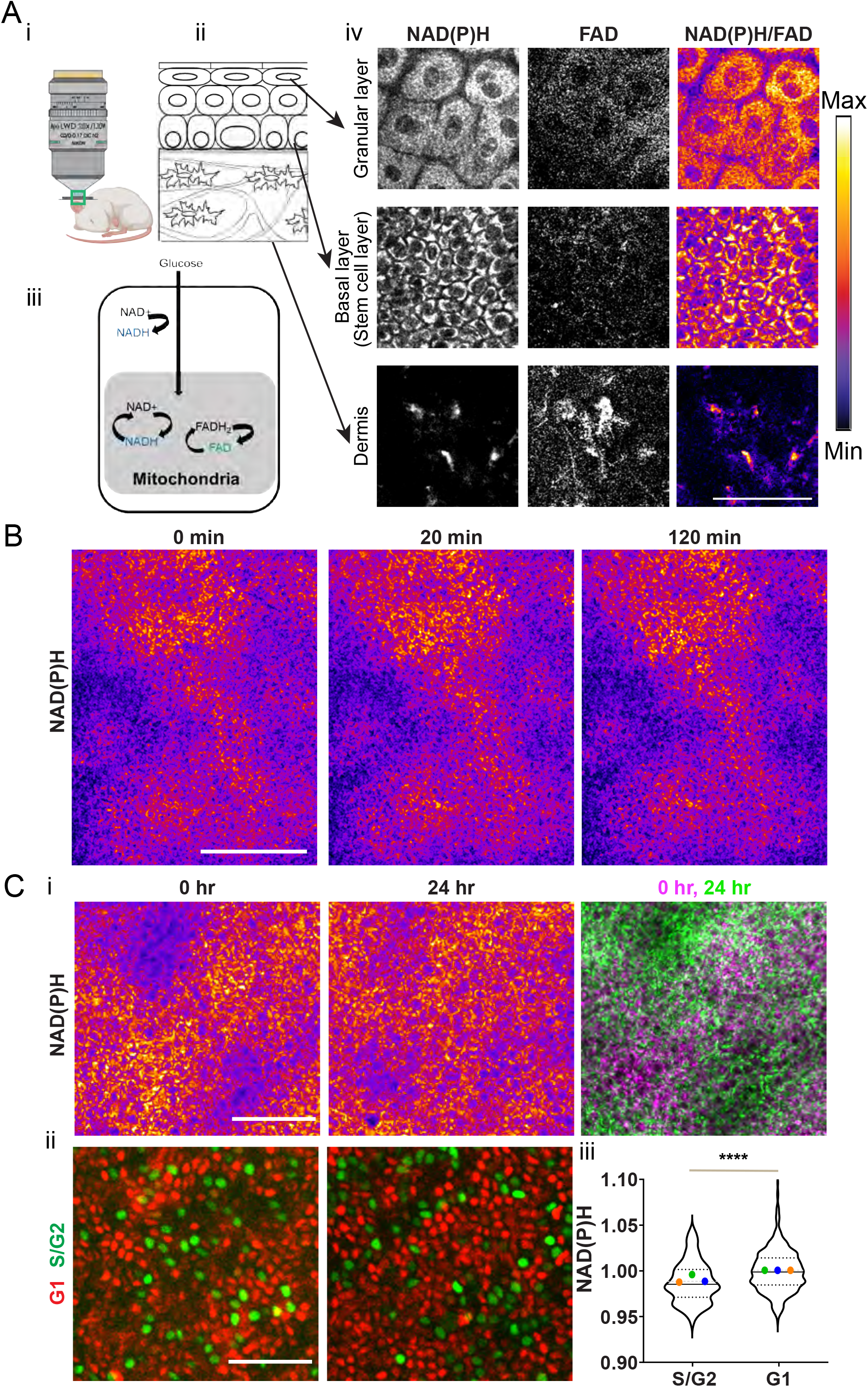
Epidermal stem cells have a unique and constrained metabolic signature at homeostasis A. **(i)** Label-free imaging of NAD(P)H and FAD in epidermis of live mice reveals **(ii)** the dynamic metabolic signature of **(iii)** the stem cell compartment. Only the reduced and oxidized counterparts NADH, NADPH, and FAD are fluorescent and together they report on the redox ratio of the cell. Changing glycolysis to mitochondrial metabolism rates will influence this ratio. **(iv)** Two-photon images of various epidermal layers and cell types in skin epithelia capture the endogenous autofluorescence of the metabolic cofactors NAD(P)H and FAD. NAD(P)H /FAD levels in the epidermal stem cell compartment (basal layer) are high compared to cells in dermis. Redox ratio (NAD(P)H/FAD) is represented as an intensity plot (scale on right). **B**. The same region of the basal epidermis is imaged over 2 hours from non-hairy epidermis (paw) to show that clusters of similar NAD(P)H intensity are stable in their spatial distribution although there are small fluctuations in intensity. **C. (i)** In contrast, the basal epidermis when revisited after 24 hours shows altered distribution of NAD(P)H intensity patterns. The underlying second harmonic signal patterns used to identify the same regions are shown in Fig. S1B. **(ii)** Cell cycle changes also occur in this timeframe. With the Fucci cell cycle reporter, green labelled nuclei represent Geminin-containing nuclei (S/G2 cell cycle stage) and red labelled nuclei represent Cdt1-containing nuclei (G1 stage). **(iii)** NAD(P)H intensities per cell measured from G1 or S/G2 nuclei show a small but significant difference in mean intensity. n=120(S/G2), 490(G1) cells from 3 mice. The average value from each mouse is labelled with a different color. All Cells (Violin plot) p-value<0.0001 (Welch’s t-test); Averages p-value<0.01 (t-test). Scale bar=50μm

Given the unique regenerative capability of epidermal stem cells, we asked whether they had a distinct metabolic signature as compared to other cell types in the dermis. We found that the cells in the epidermis have a high NAD(P)H intensity in comparison to FAD (Fig.1A iv). This is unlike many cells in the dermis, where there is a higher relative FAD to NAD(P)H intensity (Fig.1A iv; Movie S1, S2). To test if our method accurately captured NAD(P)H and FAD intensities, we treated 293T cells in culture with cyanide, which blocks the Complex IV of the electron transport chain and causes the accumulation of NADH in the cells. We observed an immediate (2-5 minutes) increase in NAD(P)H intensities and NAD(P)H/ FAD ratio (Fig S1A), consistent with previous studies, thereby confirming that our imaging parameters accurately capture the levels of changing co-metabolites that report on correspondent catabolic rates in cells.

Epidermal stem cells are constantly cycling, leading us to ask whether their metabolic signature is dynamic or stable. To address this question, we revisited the same tissue region over hours to days to track its metabolic state. In the basal stem cell layer, there were large clusters (∼25-50) of cells sharing similar NAD(P)H intensities in a pattern that remained stable over 2-3 hours of imaging (Fig.1B). However, these clusters had different relative NAD(P)H intensities (higher or lower compared to neighbors) when revisited after 24 hours (Fig.1C i). In this timeframe, the cell cycle states of many basal cells also change (Fig.1C ii). To understand how the cell cycle may influence NAD(P)H intensities, we used a reporter that distinguishes G1 cells (marked by mCherry-Ctd1) and S/G2 cells (marked by mVenus-Geminin) (Fucci 2 mice) (Fig.1C ii). We observed a small but significant difference between the mean NAD(P)H intensities of S/G2 cells and G1 cells in the basal layer, with S/G2 cells displaying a relatively lower NAD(P)H intensity than G1 cells (Fig.1C iii).

Altogether, these findings show that epidermal stem cells have a unique metabolic signature, with small variations conferred by changing cell cycle statuses in individual cells contributing to the spread of the redox values in homoeostasis.

### Redox changes precede morphological aberrancies caused by βcatGOF mutation in epidermal stem cells

We next asked how epidermal stem cell redox states change upon expression of the oncogenic mutation β-Catenin-Gain of Function (*β-catenin*^*flox(Ex3)*1,8^; referred here as βcatGOF), which is expected to affect growth rates within the tissue. Mice carrying *βcatGOF; K14CreER* were treated with Tamoxifen to activate the Cre recombinase in the basal stem cell layer (which express Keratin 14) and, in turn, induce the expression of βcatGOF in a mosaic manner. At Day 5 post-tamoxifen, the basal layer consisted of a single layer of nuclei, as in homeostasis, with increased nuclear β-catenin localization compared to control cells (Fig. 2A) as observed through immuno-staining. By Day 15, aberrant structures with several layers of closely packed nuclei enriched in nuclear β-catenin had formed and protruded into the dermis, resembling hair follicle placodes (Fig. 2A, Fig.S2A: Movie S5) (Brown, Pineda et al., 2017; Gat et al., 1998). To determine when cell behavioral changes start to emerge, we probed for proliferation events at the above time-points. Proliferation changes mirrored these morphological changes: there was no significant change in the number of phospho-Histone3 (pH3) labelled nuclei at Day 5 post-tamoxifen, but there was a significant increase in pH3-labelled nuclei at Day 15 (Fig. S2B, C). Hence, at Day 5 post-tamoxifen, even though β-catenin protein showed greater nuclear (activated) localization than in controls, we did not observe aberrancies in tissue structure or proliferation in the stem cell compartment of βcatGOF mosaic tissue (Fig. 2A, Fig.S2; Movies S3-S5).

**Figure 2:**
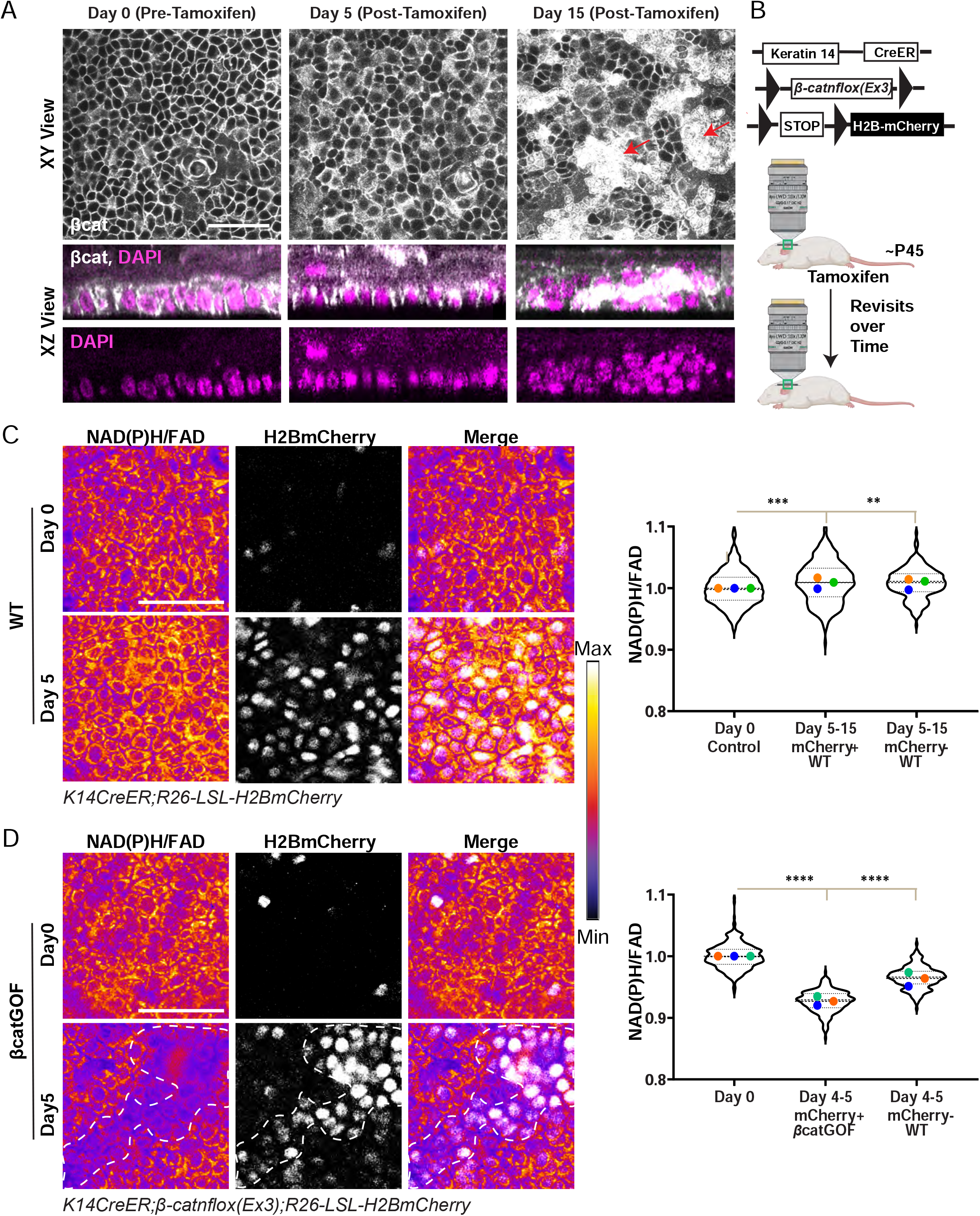
βcatGOF-induced stem cells shows rapid changes in redox (NAD(P)H/FAD) ratio before the emergence of other morphological aberrancies. **A**. β-catenin immuno-fluorescence staining in basal layer of epidermis from *K14CreER; βcatGOF* mice 0, 5, and 15 days’ post-tamoxifen-induced recombination and expression of βcatGOF. XY (top panel) and XZ (bottom panel) sections show that the 3D structure of the basal layer is unperturbed at 5 days, unlike the multiple rows of nuclei found in aberrant placodes at 15 days (eg: red arrows; More images in S2B). **B**. Schematic showing that similar regions from the same animal were imaged before and after tamoxifen-induced recombination and expression of βcatGOF and H2BmCherry (C-F). Image created with icons from Biorender. **C**. In control littermates (*K14CreER; LSL-H2BmCherry*), recombination post-tamoxifen leads to the expression of H2BmCherry (white nuclei). These recombined cells or their neighbors show no significant reduction in NAD(P)H/FAD ratio when imaged at 5, 9, or 15 days after tamoxifen administration. n(in order x-axis)=423, 250, 221 cells from 3 mice. The average value from each mouse is labelled with a different color for all graphs. Averages not significantly different. All Cells (Violin plot) **= p-value <0.003 and ***= p-value<0.0001 (One-way ANOVA; Multiple comparisons) show a slight increase (difference in means = 0.007). **D**. In βcatGOF mice (*K14CreER; βcatGOF; LSL-H2Bmcherry*), mutant cells are indicated by co-expression of nuclear H2BmCherry (white, outlined by white dashed line). Despite there being no other morphological aberrancy, the NAD(P)H/FAD ratio of mutant cells (H2Bmcherry positive) steeply drops after 5 days when compared to values at Day 0. The neighboring WT cells (H2Bmcherry negative) also show a drop in redox ratio (NAD(P)H/FAD), although to a higher range than the mutant cells. n(in order x-axis)=336, 231, 198 cells from 3 mice. Averages p-value<0.002. All Cells (Violin plot) -**** = p-value <0.0001 (One-way ANOVA; Multiple Comparisons; Day 0 vs Day 4-5 βcatGOF and WT for averages and distributions). Scale bar=50μm.

To interrogate whether metabolic changes precede or follow these morphological aberrancies in the mutant mosaic epidermis, we measured NAD(P)H and FAD levels in the stem cell layer beginning at Day 5 post-tamoxifen, prior to the appearance of aberrant phenotypes. To distinguish between and follow recombined vs. unrecombined epidermal cells (e.g. βcatGOF vs WT in the mutant mosaic epidermis) over time in the homeostatic adult mouse, we expressed *Rosa26-CAG-LSL-H2B-mCherry* (referred to as *LSL-H2BmCherry*) in the basal stem cell compartment using *K14CreER* (*K14CreER; LSL-H2BmCherry*) (Fig. 2B). First, we measured the variance of redox values in control conditions including after tamoxifen-induced recombination, expression of fluorescent alleles, and repeated imaging of the same epidermal region. We found that NAD(P)H/FAD maintains a consistent range of intensities in the homeostatic adult basal stem cell compartment in revisits 5, 9, or 15 days after tamoxifen treatment and the expression of fluorescent reporters (Fig. 2C). To understand the metabolic changes in basal cells in the context of tissue and behavioral changes that happen upon induction of the βcatGOF mutation (Fig.2A, Fig.S2A), we used Cre-induced H2BmCherry and βcatGOF co-expression to identify and track mutant βcatGOF cells over time (*K14CreER; β-catn*^*flox(Ex3)*^; *LSL-H2BmCherry*). Approximately 80% of the stem cell layer was recombined and expressed H2BmCherry and βcatGOF. Five days after induction, the βcatGOF mutant stem cells (H2BmCherry-positive) reported a rapid drop of NAD(P)H/FAD ratio (Fig. 2D). WT cells in the mosaic tissue (H2BmCherry-negative) also reported a decreased NAD(P)H/FAD ratio, but to a range distinct from and higher than that of the mutant cells (Fig. 2D), suggesting that they respond to alterations in the metabolic state of neighboring cells. Hence, the stem cell layer undergoes a rapid change in its redox state in response to the presence of βcatGOF cells before the development of morphological and behavioral aberrancies, making it one of the first observable responses to the presence of βcatGOF mutation.

### Wild-Type and βcatGOF cells in the stem cell layer show differential redox recovery over time

Over time, the βcatGOF mutation causes changes to stem cell proliferation and overall tissue architecture (Fig. 2A, S2A). Thus, we asked if the early drop in redox ratio we observed in these cells is a permanent alteration in metabolic state. By tracking the NAD(P)H and FAD intensities in the same region of basal stem cell layer over time (Fig. S3A), we observed that the cells expressing βcatGOF (H2BmCherry-positive) maintained a low NAD(P)H/FAD ratio range at 10 days post-tamoxifen, with negligible recovery from the redox ratio at 5 days post-tamoxifen (Fig. 3A,B,C). In contrast, the neighboring WT cells in the basal stem cell layer increased their NAD(P)H/FAD ratio around 2.8-fold when compared to βcatGOF cells, to resemble homeostatic redox values before induction of mutation (Day 0) (Fig. 3C). Because of this, the redox differential between the βcatGOF cells and WT cells in the stem cell layer increased over time (Fig. S3B) through the selective recovery of NAD(P)H/FAD ratios in WT cells.

**Figure 3:**
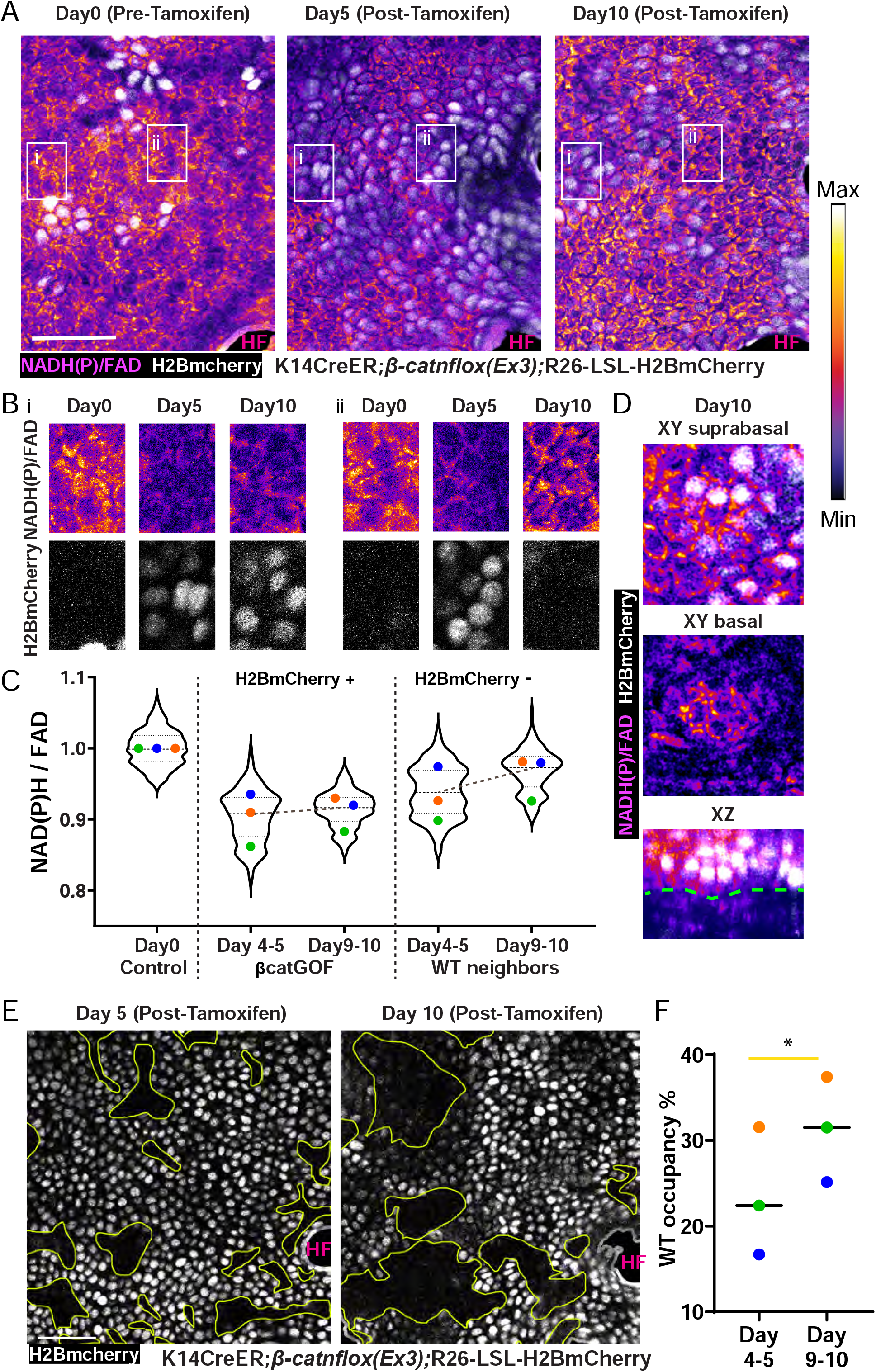
βcatGOF and WT neighbor cells have different trajectories of redox recovery over time. **A**. NAD(P)H/FAD intensities and H2BmCherry expression (white; indicates βcatGOF cells) from same region (outlined in yellow from larger regions shown in Figure S3A) in *K14CreER; βcatGOF; LSL-H2BmCherry* mice revisited 5 days and 10 days after tamoxifen administration. **B**. Insets outlined in white from panel A, highlighting regions across time wherein **(i)** βcatGOF cells at Day 5 and Day 10 show little change of redox ratio and **(ii)** βcatGOF cells (H2BmCherry positive) are replaced with WT cells (H2BmCherry negative), which have recovering redox ratios. **C**. Cells expressing βcatGOF (H2BmCherry positive) have consistently low NAD(P)H/FAD ratio at Day 4-5 and 9-11 post-tamoxifen. WT cells (H2BmCherry negative) recover their NAD(P)H/FAD ratio at Day 9-11 compared to Day 4-6. **D**. At Day 10 post-tamoxifen, the βcatGOF cells (H2BmCherry positive) are pushed to the supra-basal region, indicating differentiation (top panel), while WT cells occupy the basal layer (middle panel). Both layers can be see seen in the XZ section (bottom) with the yellow outline indicating the epidermal-dermal interface. n(in order x-axis)=379, 323, 198, 273, 227 cells from 3 mice. The average value from each mouse is labelled with a different color for all graphs. All Cells (Violin plot) p-value<0.0001 (One-Way ANOVA; Multiple Comparisons with Day0 Control). Averages p-value <0.01 for Control vs all categories except Day 9-10 WT neighbors. Day 9-10 WT neighbors not significantly different from Day 0. (RM One-Way ANOVA; Multiple measures). **E**. The βcatGOF mosaic basal stem cell layer 5 days post-tamoxifen harbors adjacent patches of WT (H2BmCherry negative; yellow outline) and mutant cells (H2BmCherry positive) which were revisited 10 days post-tamoxifen; WT patches (regions outlined in yellow) expanded between Day 5 and Day 10. F. Graph shows area occupied by WT cells from the same 300×300 μm regions from 3 mice quantified and shows in increase in coverage between Day 5 and Day 10. p-value <0.015 (Paired t-test). Scale bar=50μm.

Our previous work showed that βcatGOF cells are eliminated from outgrowths over the course of months, aided by the presence and activity of WT cells ^1^. To better understand the recovered metabolic signature of WT cells in the context of their competitive advantage, we followed the fate of the WT cells, which recovered their redox ratio, and βcatGOF cells, which did not recover their redox ratio, in the same tissue over different periods of time. We observed that between 5 and 10 days post-tamoxifen within the same region of skin, the area occupied by WT cells indicated by the absence of H2BmCherry expanded. (Fig. 3E, F). At 10 days post-tamoxifen, when the H2BmCherry-positive cells that label βcatGOF cells shrank in their coverage and WT cells expanded their coverage of the basal layer, we also observe regions where the H2BmCherry-positive mutant cells can be progressively seen enriched in the supra-basal layer, with WT cells underneath (in the basal stem cell layer) (Fig. 3D) indicating that these mutant cells are eliminated from the basal stem cell layer through differentiation. Hence, the recovery of redox in the WT cells is accompanied by their greater coverage and expansion in the stem cell layer as βcatGOF cells are eliminated from the basal stem cell layer.

While the WT and βcatGOF cells had widening redox differential shortly after induction (Fig. S3B), they also had a progressively more pronounced cell-competition phenotype when followed over a longer time span. At 1.5-2 months post-tamoxifen, there was a prominent difference in the occupancy of recombined cells in both the wild-type control littermates (*LSL-H2BmCherry;K14CreER*) and βcatGOF mosaic (*βcatGOF;LSL-H2BmCherry;K14CreER*) models. At this timepoint, the recombined cells (H2BmCherry-positive) in WT control littermates still occupied around 80% of the basal stem cell layer, in agreement with neutral drift, but in the βcatGOF mosaic epidermis, only about 25% of the basal stem cell area was occupied by H2BmCherry/βcatGOF-positive cells, reflecting a rapidly shrinking coverage by the mutant cells over time (Fig.S4A). The remaining recombined H2BmCherry/βcatGOF-positive cells were mostly packed into 3D aberrant placodes extending into the dermis (Fig. S4B) and resembling hair follicles; this also occurred in non-hairy βcatGOF mosaic skin (Fig. S4C). Altogether, this shows that WT cells recover their redox status after the initial redox drop, and ultimately outcompete the βcatGOF cells to eventually occupy more area within the basal stem cell layer.

### Hras^G12V^ cells initially decrease but ultimately recover redox state compared to WT neighbors in the basal stem cell layer

While βcatGOF mutant cells are outcompeted from the mouse epidermis by WT cells, other cell competition models include mutant cells which outcompete WT neighbors. This prompted us to ask whether the metabolic changes leading to altered redox and recovery fates that we observed in the βcatGOF mosaic are specific to the mutation itself or if they are indicative of more general cellular fitness during re-establishment of homoeostasis. To address this, we used the Hras^G12V^ (Constitutively Active Hras) mutation model where the relationship between WT and mutant cells is opposite to that in the βcatGOF model: mutant Hras^G12V^ cells outcompete WT cells in the basal stem cell layer ^1,2^ (Fig S5A). Using similar genetic backgrounds and experimental time scales as in the βcatGOF experiments described above, we asked how the NAD(P)H and FAD intensities in *K14CreER; Hras*^*G12V*^; *LSL-H2BmCherry* epidermal stem cells changed upon tamoxifen-induced recombination and expression of Hras^G12V^. We observed an initial drop in NAD(P)H and FAD intensities as well as NAD(P)H/FAD ratios at 5 days post-tamoxifen (Fig4A-C) in the basal stem cells when the same region of skin was followed over time (Fig. S5A). By following the changing NAD(P)H and FAD intensities over time points in which the recombined H2BmCherry/Hras^G12V^-positive cells expand (5 to 10 days after induction), we asked how the metabolic status of these cells changed in the context of their competitive advantage. Between 5 and 10 days post-tamoxifen, the NAD(P)H/FAD ratio of the H2BmCherry/Hras^G12V^-positive cells recovered and equalized with that of their neighboring WT cells (Fig. 4C). At 10 days post-tamoxifen, the total NAD(P)H and FAD levels in Hras^G12V^ cells had started recovering compared to Day 5 (Fig. S5 C,D). However, due to greater FAD recovery than NAD(P)H recovery at 10 days post-tamoxifen, the NAD(P)H/FAD ratio in Hras^G12V^ cells remained low (Fig. S5B), indicating higher net mitochondrial oxidation in the Hras^G12V^ cells maintained over time. Nevertheless, mutant cells in the Hras^G12V^ model recovered their total metabolite levels and redox ratios to the same values as WT neighbors (Fig. 4C). This is unlike βcatGOF mutant cells, which did not recover their total NAD(P)H levels (Fig S3C), widened their redox differential with neighboring WT cells (Fig S3B), and were ultimately out-competed by WT cells. Thus, by comparing and contrasting the “winner” Hras^G12V^ and “loser” βcatGOF mutation, we discover that both mutations initially lower the redox ratio, but the Hras^G12V^ mutation alone can recover with respect to neighboring WT cells to equalize the redox ratio across the entire basal stem cell layer.

**Figure 4.**
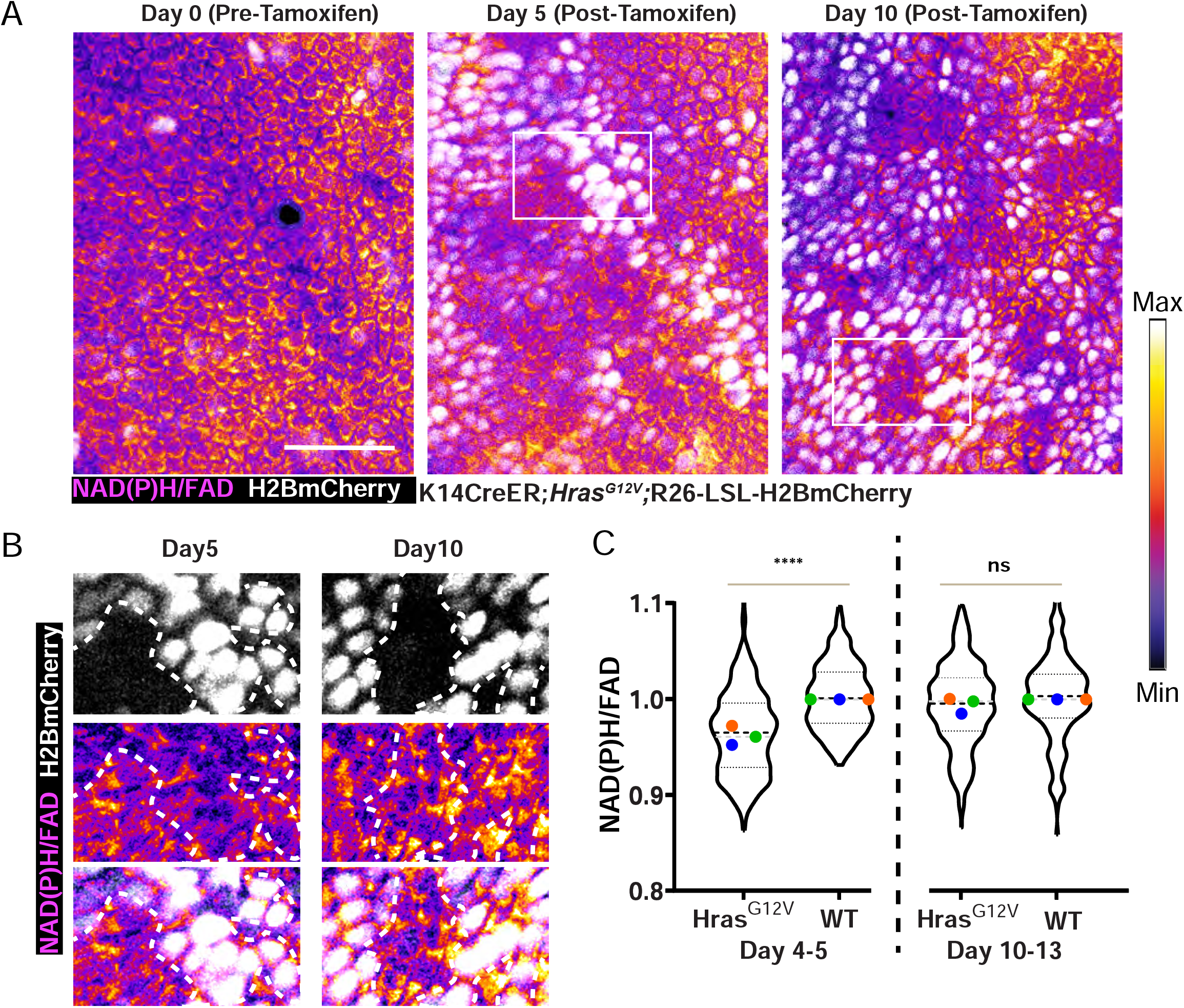
Cells with Hras^G12V^ mutation first show a drop in NAD(P)H/FAD which recovers over time. **A**. NAD(P)H/FAD intensities from same epidermal regions (outlined in yellow from larger regions shown in Figure S3D) revisited at Day 0, Days 5-6, and Days 10-13 post-tamoxifen, and expression of Hras^G12V^ and H2BmCherry (white) from *K14CreER; Hras*^*G12V*^; *LSL-H2BmCherry* mice. **B**. Insets from white outlined regions in A. Regions comprised of Hras^G12V^ and WT cells side by side show that at Day 5 (left) Hras^G12V^ (H2BmCherry positive; outlined by white dotted line) cells have a lower NAD(P)H/FAD intensity than WT cells (H2BmCherry negative). At Day 10 (right), Hras^G12V^ (H2BmCherry positive) cells have increased and recovered their NAD(P)H/FAD intensities to be similar to neighboring WT cells (H2BmCherry negative). **C**. Quantification of NAD(P)H/FAD intensities from Hras^G12V^ (H2BmCherry positive) cells normalized to their WT neighbors (H2BmCherry negative) at Day 4-5 (left) and Day 10-13 (right) to show that redox differential Day 4-5 is flattened by Day10-13. This is in contrast to βcatGOF epidermis in which the redox differential between mutant and WT increases (Fig. S3A). n (in order x-axis) =373, 180, 502, 186 cells from 3 mice. All Cells (Violin plot) **** = p-value<0.0001 (Welch’s t-test). The average value from each mouse is labelled with a different color. Average redox p-value<0.002 for Day 4-5 Hras^G12V^ vs WT; not significantly (ns) different for Day 10-13 Hras^G12V^ vs WT. Scale bar=50μm.

### Changes in glucose catabolic fluxes underlie altered NAD(P)H and FAD changes in both mutant models

NAD(P)H and FAD ratios represent the relative balance of reduced to oxidized metabolites in a cell, which in turn, depends on the relative rates of glycolysis to mitochondrial metabolism. To determine the mechanistic and biochemical basis of the changes in the NAD(P)H to FAD intensities observed through live imaging in the mutant models described so far, we asked how the glucose catabolic rates were changed in those models. We performed LC-MS/MS after infusing C_13_ isotope-labelled glucose into P45-P62 mice 6 days after injecting high doses of tamoxifen to induce expression of βcatGOF or Hras^G12V^ in most if not all epidermal cells. Epidermal tissue isolated from the mice infused with ^13^C_6_ glucose (Methods) was then used to assess the label retention at various steps of glucose metabolism to determine the flux through various steps (Fig. 5A)^3^. We found that rates of glucose flux through the TCA cycle as indicated by V_PDH_/V_CS_ (ratio of pyruvate dehydrogenase flux to citrate synthase flux, i.e. the fraction of the TCA cycle fueled by glucose) ^3^ were upregulated in both βcatGOF and Hras^G12V^ epidermis. The TCA cycle was fueled almost entirely by glucose in both mutant tissues (Fig. 5B). C_13_-labelled CO_2_ relative to glucose was also upregulated in both mutant tissues (Fig. 5C), indicating that flux through the TCA cycle was enhanced by the mutations. This is contrary to the expectation that oncogenic mutations would lead to upregulated glycolysis at the expense of downstream steps, according to the Warburg effect ^21^. Interestingly, this enhanced rate of flux through the TCA cycle is in agreement with the reduction in NAD(P)H to FAD ratio (reduced to oxidized metabolite ratio) that we observed through live imaging (Fig2C-D, 4A-C), providing a direct parallel biochemical validation.

**Figure 5:**
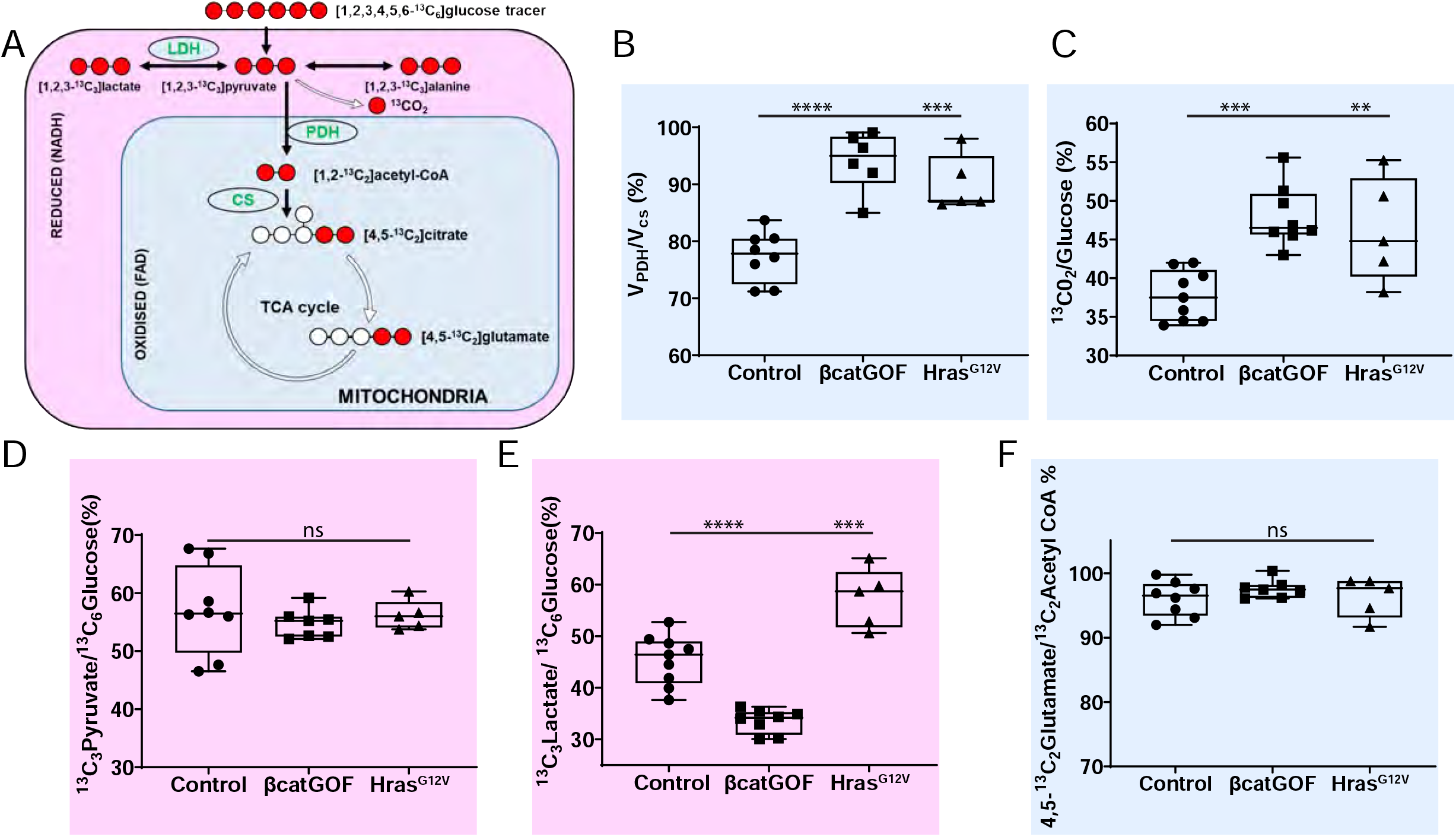
Changes in glucose catabolic fluxes underlie altered NAD(P)H and FAD changes in mutant conditions. **A**. Schematic showing the labelling of carbon (red) in downstream metabolites when mice are infused with ^13^C_6_ glucose. The readouts which pertain to reduced NADH in the cytosol are labelled in pink and those pertaining to mitochondrial metabolism are labelled in blue. **B**. V_PDH_/V_CS_ ratio represents the contribution of glucose to TCA flux in the epidermis. In both βcatGOF and Hras^G12V^ epidermis, V_PDH_/V_CS_ increases, indicating accelerated flux through the TCA cycle. **C**. Another measure of TCA flux, ^13^CO_2_ normalized to ^13^C_6_ glucose, also increases in both mutants. **D**. In contrast, labelled pyruvate normalized to labelled glucose is unchanged in both mutants. **E**. Lactate, which is considered the end-point of glycolysis, is decreased in βcatGOF while in the Hras^G12V^ epidermis, the ^13^C_3_ labelled lactate to ^13^C_6_ glucose ratio is increased. Hence the mutant Hras^G12V^ epidermis has an upregulated pyruvate to lactate flux when compared to the βcatGOF tissue, in which the pyruvate to lactate flux is downregulated. **F**. ^13^C_2_ labelled Glutamate to Acetyl CoA is a measure of the dilution of the label due to glutamine entry into the TCA cycle. The nearly 100% ratios suggest that there is not much entry of glutamine. This is not significantly different in either mutant epidermis when compared to WT epidermis. p-value **** = <0.0001, *** = <0.0005, ** = <0.0058 (One-way ANOVA, Multiple comparisons with respect to Control). Each point represents epidermis (ear) from one mouse.

To understand whether these changes in TCA flux were also reflected in upstream glycolytic steps, we asked whether the rates of glycolysis were changed. Specifically, we focused on pyruvate and lactate labelling, which is expected to be higher in oncogenic mutations according to Warburg effect ^21^. At early steps of glycolysis, specifically the transfer of ^13^C label from glucose to lactate, the two mutations differed in their effects: cells in the Hras^G12V^ epidermis upregulated flux to lactate while cells in the βcatGOF epidermis downregulated it (Fig. 5E). However, there was no parallel upregulation of glucose to pyruvate flux in either mutant epidermis (Fig. 5D), pointing towards a specific and differential modulation of pyruvate to lactate flux in the two mutants when compared to control epidermis. On the other hand, the ^13^C labelled glutamate derived from ^13^C Acetyl CoA (m+2 C_4_C_5_ glutamate/m+2 acetyl CoA) was not different from the ratio of labelled glutamate to pyruvate in the two mutant models, indicating that the glutamine contribution to the TCA cycle (i.e. dilution of glutamate) was minimal and did not differ between the two mutant models (Fig. 5F).

Altogether, we uncovered the glucose catabolic basis that drives the initial reduction in redox ratios captured through live imaging in βcatGOF and Hras^G12V^, while also revealing differences in upstream glycolytic rate in the two mutant tissues. Thus, we revealed the nodes of regulation unique to the winner and loser mutations that define their different redox behaviors.

## Discussion

With unprecedented resolution of metabolic state and ability to follow cell behaviors over time in a complex tissue-scale process like cell competition *in vivo*, we discover that the NAD(P)H/FAD ratio is cell-type specific and rapidly lowers in response to oncogenic mutations, prior to any observable cellular morphological and behavioral changes. The metabolic changes thus emerge as a first line of response to oncogenic mutations. The reduction in NAD(P)H to FAD ratio following the induction of oncogenic mutations suggests increased net mitochondrial oxidation, and we show through glucose flux measurements that this is caused by an increase in flux of glucose through TCA cycle downstream of glycolysis. However, our glucose flux measurements also reveal a differential change in tissue carrying the “winner” mutation, Hras^G12V^, wherein the flux from pyruvate to lactate is upregulated. In contrast, pyruvate to lactate flux is reduced in tissue carrying the “loser” mutation, βcatGOF. Over time, redox ratio recovers selectively in the population of stem cells that acquire a “winner” fate. In the βcatGOF model, WT cells recover their redox ratio and outcompete the mutant cells to occupy most of the basal stem cell compartment of skin and “win”. In contrast, in the Hras^G12V^ model, the mutant cells regain their redox values compared to their WT neighbors, outcompete the WT cells, and become the “winner” (Fig. 6). Hence, recovery of redox and selective upregulation of glycolytic steps, uncoupled from TCA cycle, is a hallmark or signature that predicts the ability of the cell to survive in the epidermal stem cell compartment and adopt a “winner” fate.

**Figure 6:**
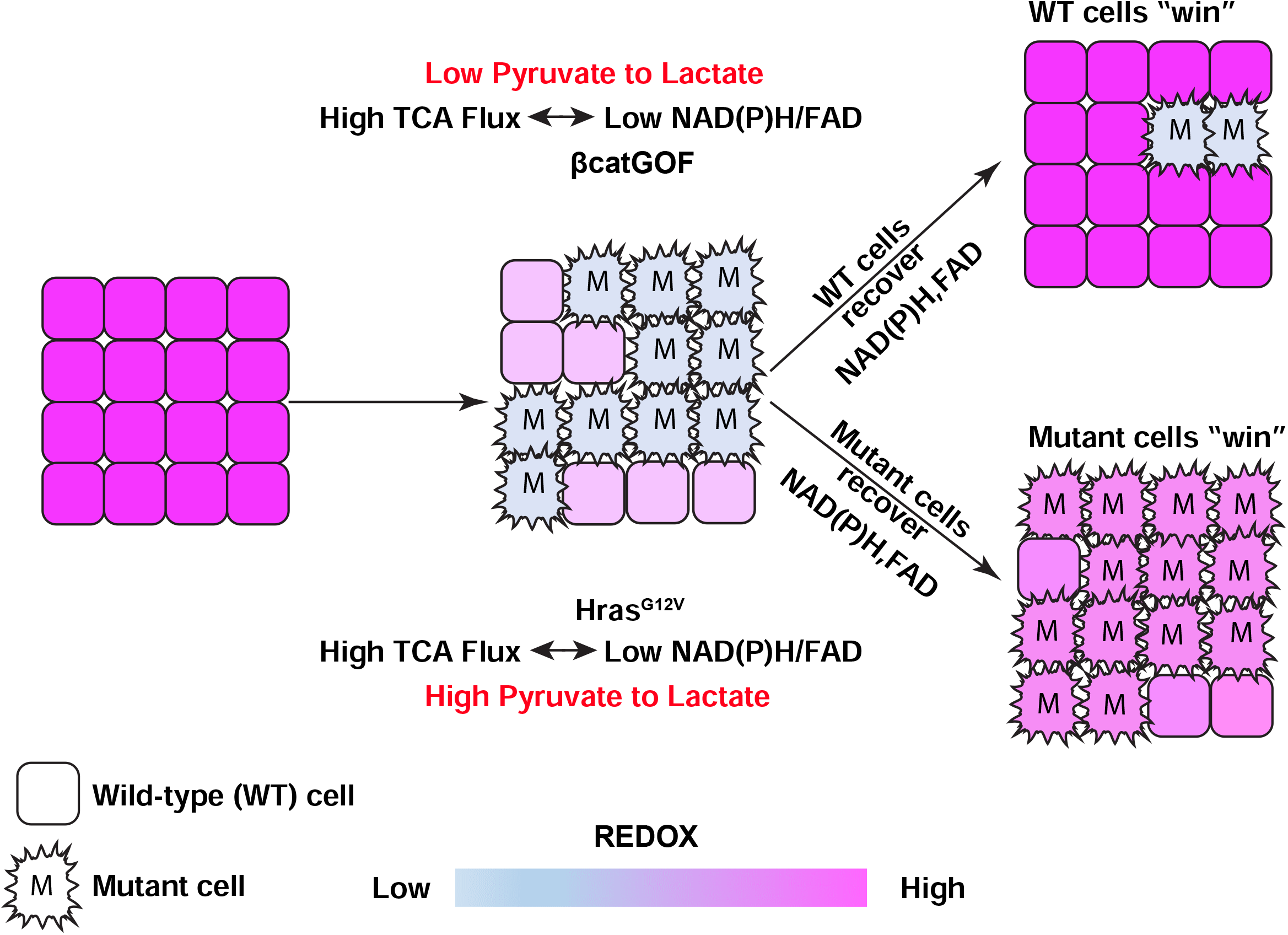
Model: Unique metabolic hallmarks of redox recovery and glycolytic rates identify the winner cells. NAD(P)H/FAD (reduced to oxidized metabolites) maintains a robust range in the homeostatic skin epidermal stem cells as measured through optical redox ratio imaging. Upon induction of the mutation in the basal layer of the epidermis in a mosaic manner, there is a rapid reduction in NAD(P)H/FAD in both the mutant and WT cells, at early time-points preceding morphological aberrancy. In parallel, as glucose tracer-assisted mass spectrometry analysis shows, the mutant tissue also accelerates its TCA flux in line with the reduction in NAD(P)H/FAD ratio. Redox imaging of the mutant epidermis and revisits over time reveals that the redox recovery is limited to winner cells. In the βcatGOF mutant model, the winner wild-type (WT) cells neighboring the mutant cells recover their NAD(P)H/ FAD ratios whereas in the Hras^G12V^ mutant model, the winner mutant cells recover and equilibrate their redox ratios with their neighboring WT cells. These redox changes happen while the loser βcatGOF mutant cells are outcompeted by WT cells and the winner Hras^G12V^ cells outcompete the WT cells in the basal stem cell layer of the epidermis. The winner and loser mutant cells also have a marked difference in glycolytic rates, wherein the winner Hras^G12V^ cells upregulate pyruvate to lactate flux while this step is downregulated in the βcatGOF mutant epidermis. Thus, the winner and loser mutant cells have unique redox recovery trajectories and glycolytic rates contributing to their identity and eventual cell-competition outcomes.

The direct observation of NAD(P)H and FAD levels with genetic and cellular resolution *in vivo*, coupled with direct measurements of glycolytic and TCA cycle fluxes enables an accurate readout of the tissue metabolic state. From these studies, a unique picture of the epidermis emerges wherein a high NAD(P)H to FAD ratio implies high glycolytic activity, yet the high V_PDH_/V_CS_ also shows that glucose fuels up to 80% of the TCA cycle in these cells. This high reliance of the TCA cycle on glucose is not typical of most other tissues like heart, liver, kidney, and muscle which show a far lower V_PDH_/V_CS_ between 20 and 40% ^3^. It remains to be seen whether this high glycolytic rate and glucose oxidation are a characteristic of rapidly cycling tissues with an active stem cell pool. Interestingly, the brain also exhibits a high V_PDH_/V_CS 3_, similar to skin epidermis, raising the possibility that their developmental origin (ectoderm derived) could also define these metabolic signatures. Strikingly, in the presence of the oncogenic mutations βcatGOF and Hras^G12V^, glucose oxidation increases and almost reaches 100%. Although this is paradoxical to the conventional understanding of the Warburg effect, an increasing number of *in vivo* studies suggest that increased glucose uptake and glycolysis does not always come at the expense of the TCA cycle and mitochondrial oxidation ^33,37–39^. In fact, these pathways may have a cooperative role in supporting proliferation and cell survival. In our study, the “winner” mutation Hras^G12V^, in addition to increased glucose cycling through TCA cycle and hence oxidation, also upregulates flux to lactate. This ability to upregulate and modulate both the glycolytic and glucose oxidation pathways could be an adaptation that characterizes cell survival during cell competition.

Enhanced pyruvate to lactate flux (considered the glycolytic end product) could support a sustained increase in mitochondrial oxidation in the Hras^G12V^ mutant epidermis by regenerating the NAD+ precursor. Although pyruvate to lactate flux through NAD+ regeneration has been proposed as a mode of maintaining redox balance in cells with enhanced glycolysis ^16^, in the Hras^G12V^ mutant model the regeneration of NAD+ could help sustain the enhanced flux through the TCA cycle and recover the NADH and FAD (Fig. S5 C, D) over time. This is unlike in βcatGOF cells where the lower NAD(P)H to FAD ratio is at the expense of total NAD(P)H levels (Fig. S3C), perhaps becoming unsustainable over time and contributing to poor cellular fitness when compared to neighbors. Alternately, lactate itself could directly contribute to the “winner” cell fate in Hras^G12V^ if it was secreted, potentially communicating redox state to neighboring WT cells and enabling rapid equilibration of redox state between Hras^G12V^ and WT neighbors (Fig. 4C). Such cell-to-cell lactate transport has been implicated in the maintenance of homoeostasis and cell survival in other contexts like the astrocyte-neuron lactate shuttle ^40^ and survival of hypoxic tumor cells ^41,42^. Furthermore, Lactate dehydrogenase (LDH) itself, which catalyzes the conversion of pyruvate to lactate, as well as lactate transporters (Monocarboxylate Transporters) have critical functions in stem cell activation in the hair follicle^43^. LDH could be differentially activated in the βcatGOF and Hras^G12V^ cells, perhaps through differential activation of c-Myc (directly downstream of Hras), which is known to upregulate LDH activity. Thus similar to other models ^44,45^, lactate flux emerges as a central node of regulation and difference between the “winner” and “loser” mutations in the skin.

Cell-cell communication of metabolic state across the epidermal basal stem cell layer is supported by three lines of evidence. First, the presence of mutant βcatGOF lowers the redox ratio of neighboring WT cells (Fig. 2D). Interestingly, the effect of βcatGOF cells on the metabolic state of WT cells progressively decreases as these cells are eliminated from the 2-D basal stem cell layer (Fig. S4B) even though some of these mutant cells are maintained within as placodes resembling hair follicles in both hairy and non-hairy skin (Fig. S4). Secondly, Hras^G12V^ cells can regain and equalize their redox ratio with their WT neighbors (Fig. 4C) through the modulation of NAD(P)H and FAD values between 5 and 10 days after Hras^G12V^ induction. This suggests a role for cell-cell junctions in mediating the communication of redox state. Third, the homeostatic basal layer is composed of clusters of cells with similar NAD(P)H values that are stable over hours and dynamic in their position over days in tune with the turnover of cell cycle in these cells (Fig.1 B-C). Exploring the composition and nature of this cell-cell communication as well as the metabolite(s) and carrier(s) that communicate the redox state of a cell to its neighbor would be interesting avenues for further studies. As discussed above, lactate transport is one such candidate mechanism of communication.

The language of cell competition – “winner”, “loser”, “fitness” – implies the existence of an ideal state, in this case, an ideal redox ratio that characterizes the cells of a tissue. While we do uncover metabolic signatures that are maintained during homeostasis in the tissue, our study also shows that winners and losers are context dependent. WT cells are losers when compared to Hras^G12V^ neighbors yet modulate their redox ratios in the presence of βcatGOF neighbors and ultimately outcompete them. Thus, rather than an ideal metabolic state, metabolic plasticity plays an important role in the maintenance of homoeostasis in the pre-cancerous epidermis. The changes in NAD(P)H levels, FAD levels, and glucose catabolic flux precede changes in proliferation and morphological aberrancies. This implies that manipulating redox state can modulate the cell competition outcome, which has profound implications for therapeutically eliminating oncogenic mutations from the skin epidermis.

## Acknowledgements

We also thank all members of the Greco lab, especially Tianchi Xin for critical feedback on the manuscript as well as advice on technique and tool development. Funding: This work is supported by an HHMI Scholar award and NIH grants number 1R01AR063663-01, 1R01AR067755-01A1, DP1AG066590-01, R01AR072668 (all to V.G), and 1R37CA258261-01 (to R.J.P).

## Author Contributions

A.H., R.J.P. and V.G. designed experiments. A.H. performed 2-photon imaging, mouse genetics, and image analysis. Z.L, R.J.P and A.H. performed mass spectrometric studies. M.S, D.G and D.G.G. advised and assisted with optimizing redox imaging within our *in vivo* imaging platform. D.G.G. assisted with development of analysis tools and analysis. E.L. and K.T. performed whole mount staining and image analysis. C.M did FACS studies and S.G. assisted with mouse genetics. A.H., R.J.P. and V.G. wrote the manuscript with input throughout from L.G.

---

**Supplementary Figure 1:**
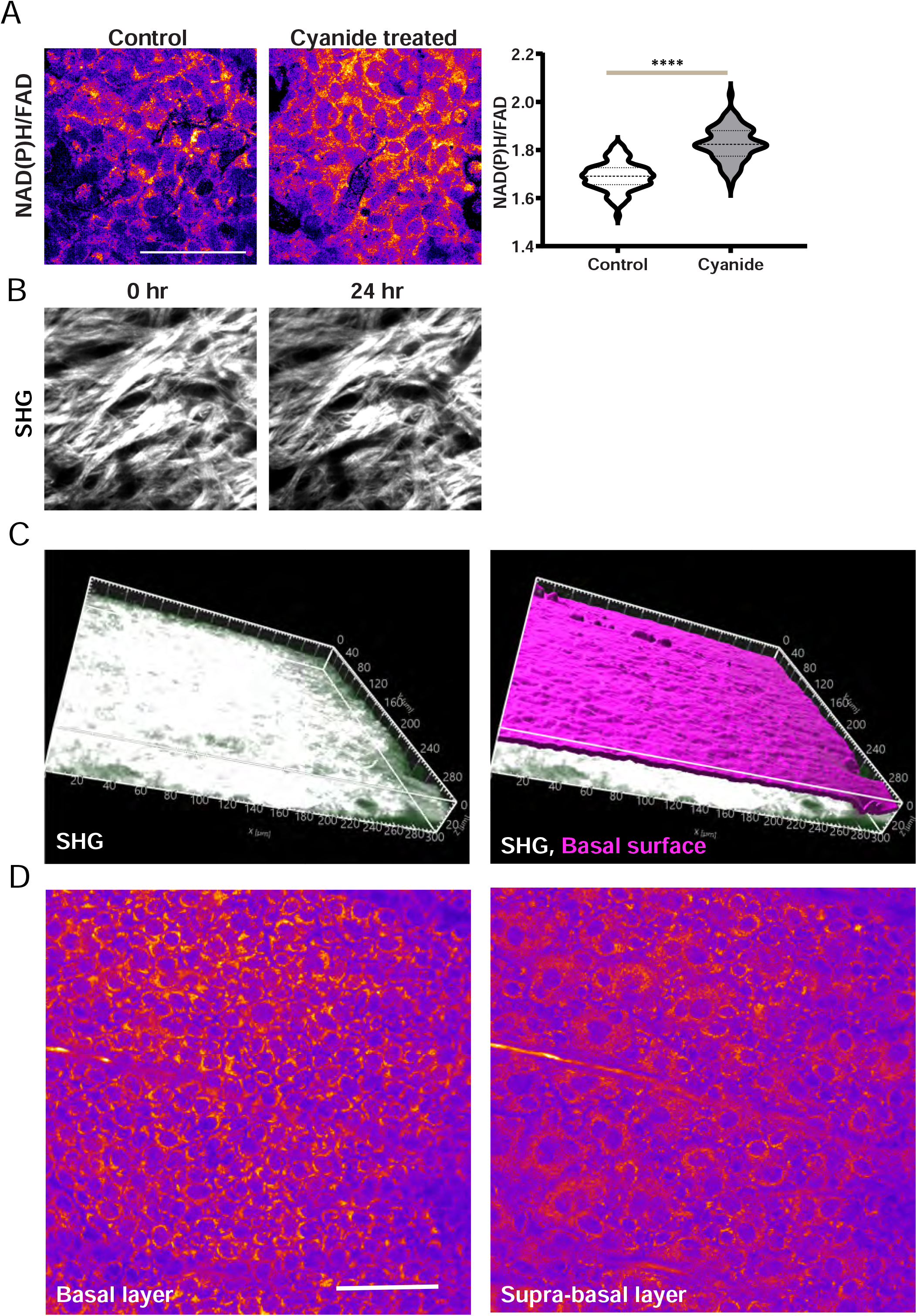
Imaging NAD(P)H and FAD. **A**. NAD(P)H/FAD intensities from HEK293T cells treated with 4mM Sodium cyanide imaged after 5 minutes of adding cyanide. NAD(P)H/FAD ratio rapidly increases in response to the inhibition of Complex IV and oxidative phosphorylation, as expected, because blockage of mitochondrial electron transport leads to accumulation of NAD(P)H in the cells. **** = p-value <0.0001(t-test). n(In order x-axis) = 86, 78 cells, representative of 2 independent experimental replicates. **B**. The second harmonic signal (SHG) from collagen in regions from Fig. 1C and 1D at 0 hours and revisited after 24 hours. The collagen fibrils and blood vessels are used to identify the same region of epidermis. **C**. 3-D projection from the Imaris software used to isolate the basal layer. After thresholding and surfacing the second harmonic signal (SHG) using the Distance Transform function in Imaris, pixels from selected distances from the epidermal-dermal interface are isolated and projected to isolate the basal layer. **D**. NAD(P)H from Basal (left) and supra-basal (right) layers isolated at different distances from the epidermal-dermal interface as described in C. Scale bar=50μm.

**Supplementary Figure 2:**
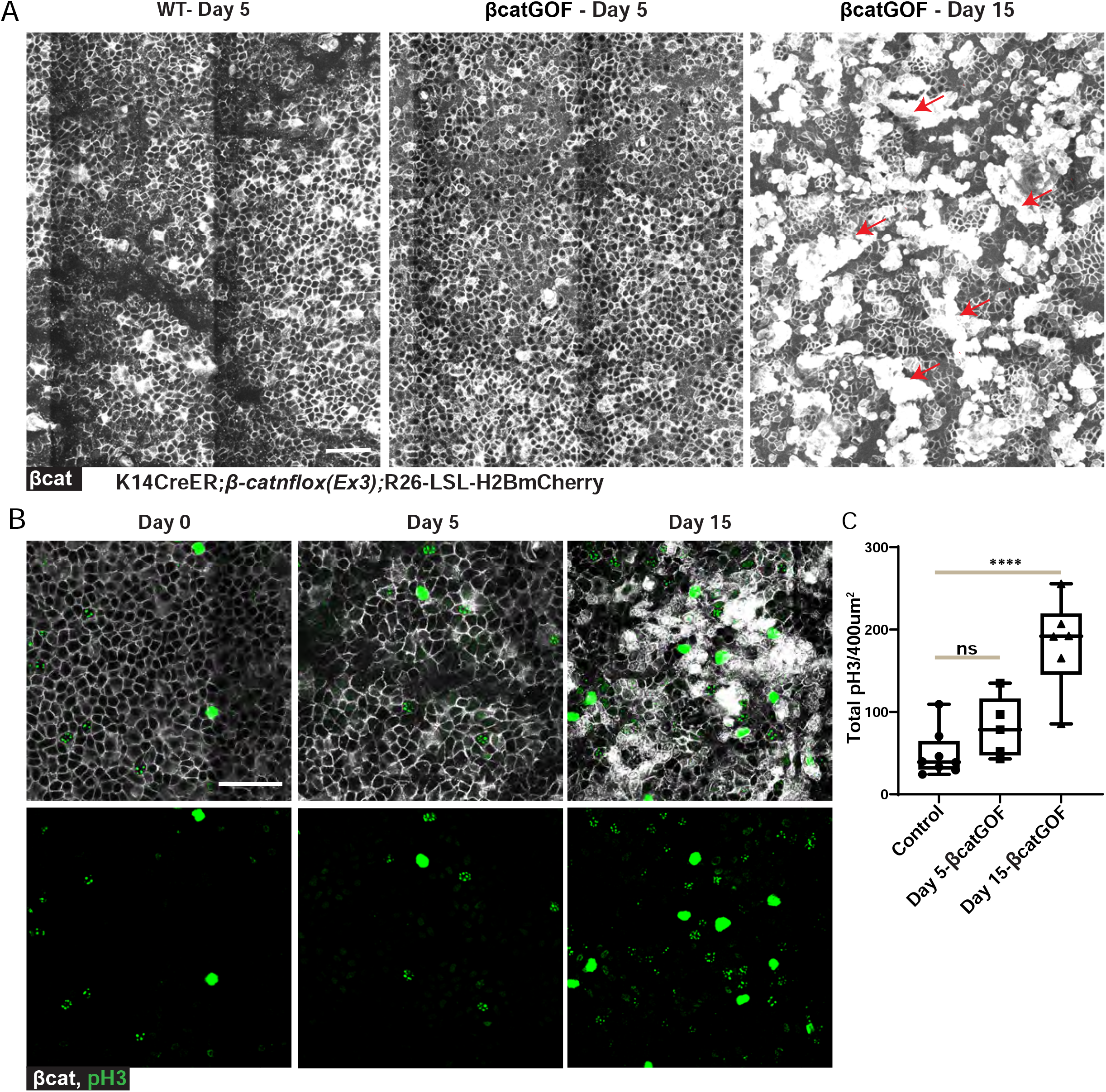
Morphological and Behavioral changes in the βcatGOF mutant epidermis. **A**. β-catenin immunofluorescence staining (grayscale) in the basal layer of epidermis from large regions in mice of genotype *K14CreER; βcatGOF* 5 and 15 days’ post-tamoxifen-induced recombination and expression of βcatGOF, similar to Fig 2A. Control shown is from littermate without mutation 5 days post-tamoxifen. The morphology of the basal layer is indistinguishable between Day 0 and Day 5. However, at Day 15, several aberrant structures (placodes) are visible (examples: red arrows) in the basal layer with stacked layers of nuclei enriched in β-catenin. More magnified images of these regions are in Fig 2A. **B**. phosho-Histone3 (pH3-green) and β-catenin staining in epidermis at Day 0 (before tamoxifen), Day 5, and Day 10 post-tamoxifen. **C**. pH3 positive nuclei were counted and quantified from 400×400 μm^2^ regions (each point on the graph) from 3 mice. **** = p-value<0.0001, ns = not significant (One-way ANOVA; Multiple comparisons with respect to Control/Day0). Scale bar= 50μm

**Supplementary Figure 3:**
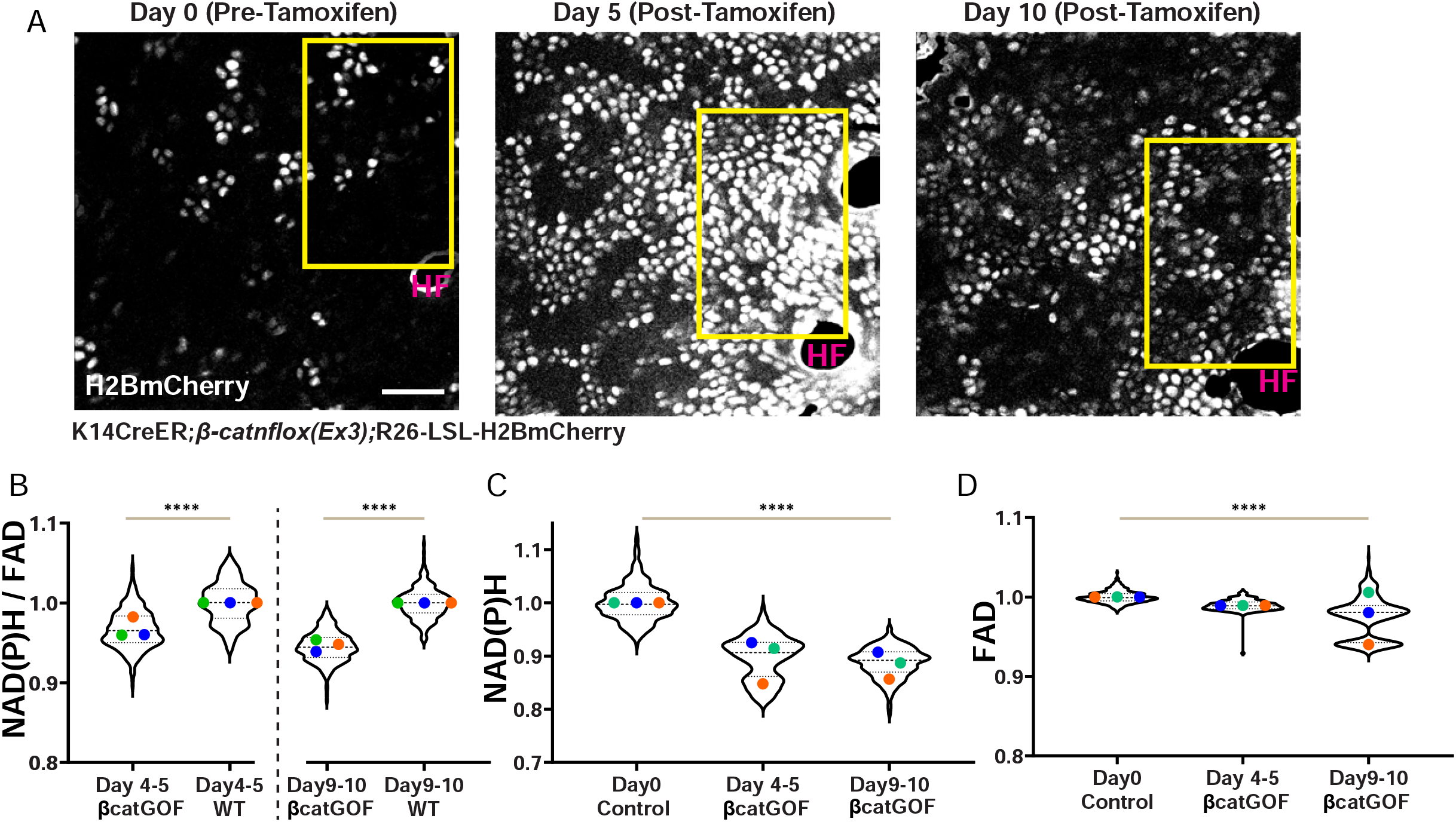
Revisits of the same epidermal regions to show different cell-competition phenotypes and redox changes in βcatGOF mutant epidermis. Morphological hallmarks including blood vessels, hair follicles (labelled as HF), and second harmonic signal are used to revisit the same region in the mouse ear epidermis over many days (Methods). **A**. The larger regions from which the epidermal revisits magnified in Fig 3A (outlined in yellow) are shown. White nuclei (H2BmCherry positive) label the cells that are recombined and hence are considered βcatGOF mutant in mice of genotype *K14CreER; βcatGOF; LSL-H2BmCherry*. **B**. The NAD(P)H/FAD per cell values from βcatGOF cells (H2BmCherry positive) and neighboring WT cells (H2BmCherry negative) at Day 4-6 and Day 9-11 post-tamoxifen shows that redox differential between βcatGOF cell and WT cells increases between 5 and 10 days because of the selective recovery of WT cells. Data shown in Fig 3C are replotted here to show that βcatGOF mutant cells increase their redox differential in contrast to Hras^G12V^ epidermis (Fig 4C), where the redox difference is flattened. n (in order x-axis) =323, 198, 273, 227 cells. All Cells (Violin plot) p-value **** = <0.0001 (Welch’s t-test). Averages p-value<0.01 for Day 4-5; p-value<0.002 for Day 9-10 (t-test). **C-D**. Total NAD(P)H and FAD from βcatGOF-induced epidermis shows no recovery of NAD(P)H and FAD values in the βcatGOF cells between 5 and 10 days’ post-tamoxifen induction. n(in order x-axis)=379, 323, 273. All Cells (Violin plot) p-value<0.0001 (One-Way ANOVA). NAD(P)H Averages p-value<0.02. FAD averages not significantly different (One-Way ANOVA). Scale bar= 50μm.

**Supplementary Figure 4:**
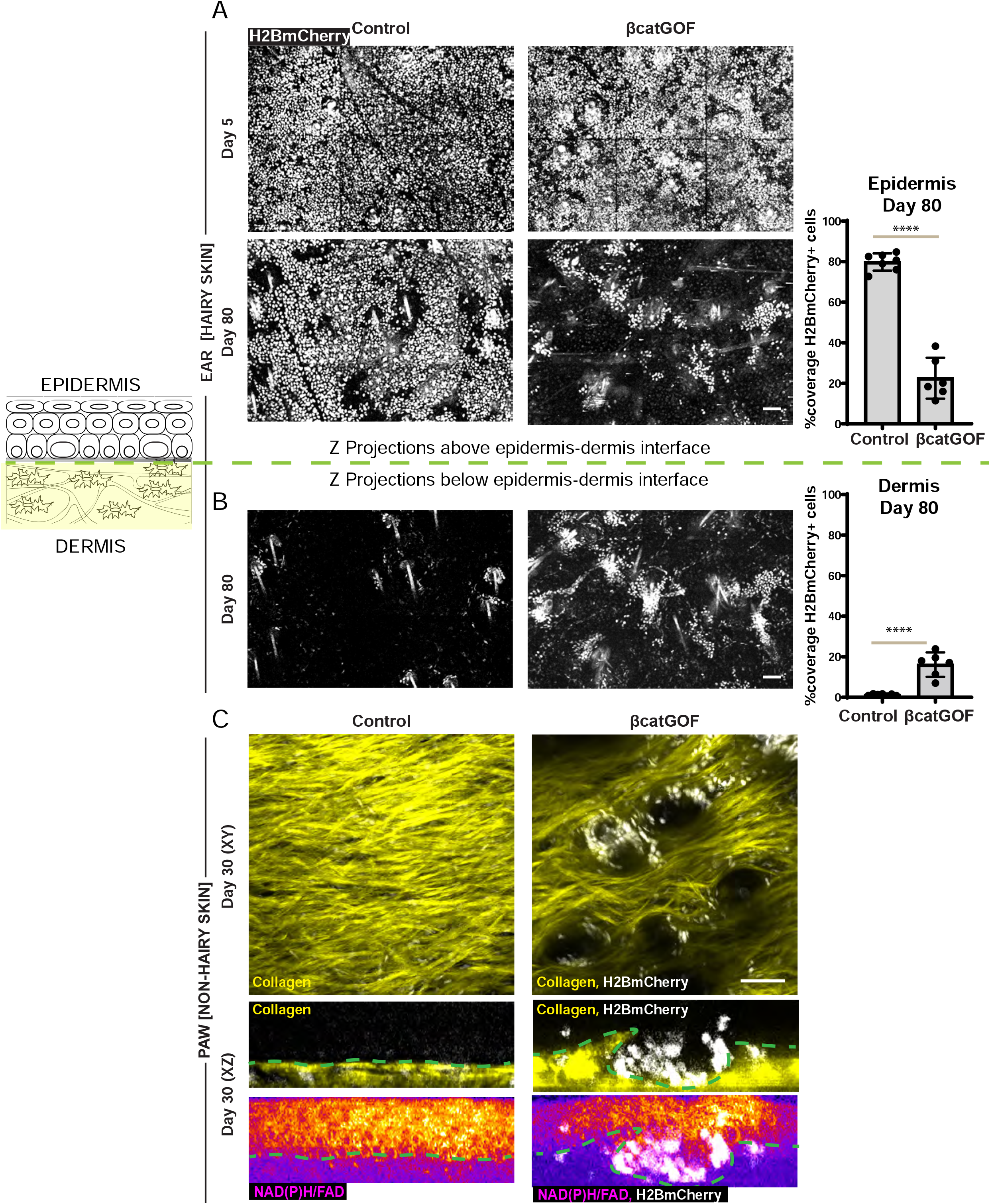
WT cells outcompete βcatGOF after 1-2 months while the majority of the remaining βcatGOF cells are found in placodes that extend into the dermis. **A**. Z-projections from large regions of skin above the epidermal-dermal interface from Control (genotype: *K14CreER; LSL-H2BmCherry*) and βcatGOF-induced mice (genotype: *K14CreER; βcatGOF; LSL-H2BmCherry*) shows similar rates of recombination, with recombined cells (mCherry+; white nuclei) representing control (left) or mutant cells (right) 5 days after recombination covering ∼80% of the epidermis five days post-tamoxifen. However, after two months (bottom panel), while control tissue retains ∼80% (graph right), βcatGOF tissue has reduced the coverage of mutant cells from ∼80% to ∼25%. **B**. The majority of recombined mutant cells (H2BmCherry positive; white nuclei) extend in sub-epidermal structures taking up ∼20 % of the sub-epidermal space compared to negligible coverage of H2Bmcherry positive epidermal cells in controls. ****= p-value< 0.0001 (t-test) from ∼1200-1400 μm^2^ regions (each point) from the ears of 2 mice. **C**. In non-hairy skin (paw), the βcatGOF-induced cells (H2BmCherry positive; white nuclei) are found in long-lived hair follicle-like structures in the βcatGOF-induced epidermis (right). In contrast, in control tissue (*K14CreER*) the second harmonic signal (SHG) from collagen in dermis (yellow) is uninterrupted by hair follicles. XZ sections show the placode-like structures with βcatGOF cells that extend into the dermis with an altered NAD(P)H signal that interrupts the collagen (SHG). Scale bar=50μm.

**Supplementary Figure 5:**
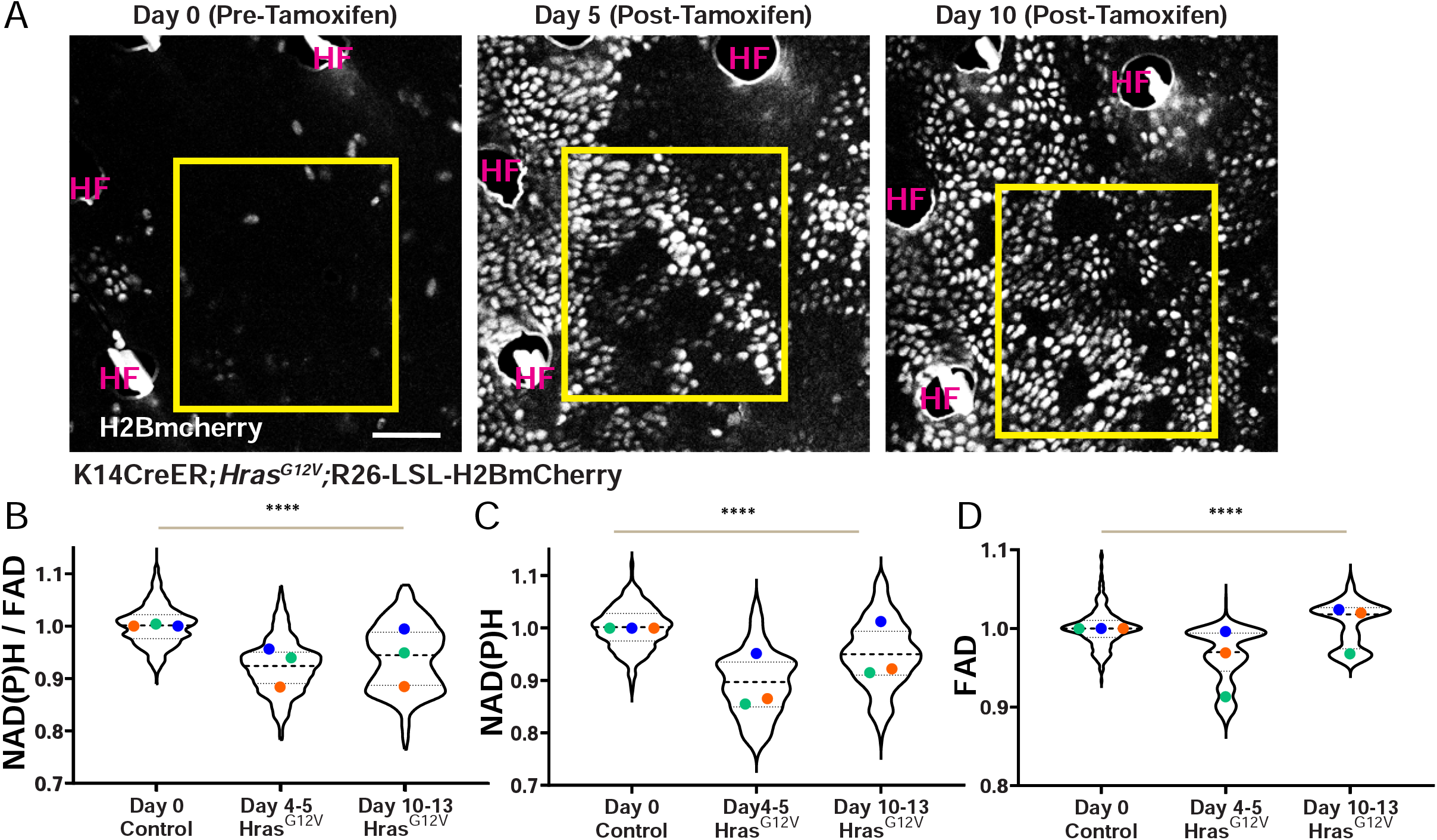
Revisits of the same epidermal regions shows different cell-competition phenotypes and redox changes in the Hras_G12V_ mutant model. **A**. White nuclei (H2BmCherry positive) label cells that are recombined and carry the Hras^G12V^ mutation in mice of genotype *K14CreER; Hras*^*G12V*^; *LSL-H2BmCherry*. **B, C, D**. NAD(P)H/FAD (B), NAD(P)H (C) and FAD (D) intensities from Hras^G12V^ cells within the epidermis revisited before tamoxifen (Day 0), Day 4-5, and Day 10-13 post-tamoxifen. The reduction in the redox distributions of NAD(P)H, FAD, and NADH(P)H/FAD intensities at Day 4-5 begins to recover at Day 10-13. Because of a greater relative FAD recovery when compared to NAD(P)H, there is only a modest recovery in the distribution of NAD(P)H/FAD intensities when compared to Day 4-5. n(in order x-axis) = 378, 409, 431 cells. All Cells (Violin plot) p-value< 0.0001 (One-Way ANOVA with respect to Day 0) shown in B-D. Averages: Day 0 Control vs Day 4-5 Hras^G12V^ p-value <0.02 for NAD(P)H and NAD(P)H/FAD (t-test). Others not significant.

**Supplementary Video Movie S1: Epidermal cells have a unique metabolic signature at homeostasis:** Movie showing z stack through different layers of the skin in a live mouse. As shown in the illustration (left) NAD(P)H/ FAD intensities are represented in a fire lut scale from the top differentiated cells to the basal stem cell layer of the epidermis followed by different cell types in the dermis. Collagen fibers in white indicate the planes where the dermis starts. Scale bar=10um

**Supplementary Video Movie S2: NAD(P)H and FAD through the different layers of skin:** The NAD(P)H(left) and FAD(middle) channels and merged(right) separated from the z stack shown in Movie S1 through the different layers. from the top differentiated cells to the basal stem cell layer of the epidermis followed by different cell types in the dermis. Collagen fibers in yellow indicate the planes where the dermis starts. Scale bar= 10um.

**Supplementary Movie S3: β-catenin staining in the epidermis**. Movie showing z stack through β-catenin immuno-fluorescence (white) stained epidermis from *K14CreER; βcatGOF* mice before tamoxifen (Fig.2A). Nuceli stained with DAPI shown in magenta. Scale bar= 10um.

**Supplementary Movie S4: βcatGOF induced epidermis does not show morphological aberrancy at Day5:** Movie showing z stack through β-catenin immuno-fluorescence (white) stained epidermis from *K14CreER; βcatGOF* mice five days’ post-tamoxifen induced recombination and expression of βcatGOF. Although the β-catenin is more diffused through the cytosol, the basal layer shows stereotypical organization without any aberrance. Scale bar= 10um.

**Supplementary Movie S5: βcatGOF induced epidermis develops aberrant structures by Day 15:** Movie showing z stack through β-catenin immuno-fluorescence (white) stained epidermis from *K14CreER; βcatGOF* mice 15 days’ post-tamoxifen induced recombination and expression of βcatGOF. Aberrant placodes or outgrowths with closely packed layers of nuclei (DAPI; magenta) seen only at Day15. Scale bar= 10um.

